# *Plasmodium falciparum* increases its investment in gametocytes in the wet season in asymptomatic individuals

**DOI:** 10.1101/2020.08.04.236950

**Authors:** Colins O. Oduma, Sidney Ogolla, Harrysone Atieli, Bartholomew N. Ondigo, Ming-Chieh Lee, Andrew K. Githeko, Arlene E. Dent, James W. Kazura, Guiyun Yan, Cristian Koepfli

**Author notes:** Correspondence, Department of Biological Sciences & Eck Institute for Global Health, 319 Galvin Life Sciences, University of Notre Dame, Notre Dame, IN, 46556-0369, USA.

## Abstract

In many regions, malaria transmission is seasonal, but it is not well understood whether *P. falciparum* modulates its investment in transmission in response to seasonal vector abundance. In two sites in western Kenya (Chulaimbo and Homa Bay), we sampled 1116 asymptomatic individuals in the wet season, when vectors are abundant, and 1743 in the dry season. We screened for *P. falciparum* by qPCR, and gametocytes by *pfs25* RT-qPCR. Parasite prevalence in Chulaimbo and Homa Bay was 27.1% and 9.4% in the dry season, and 48.2% and 7.8% in the wet season respectively. Mean parasite densities did not differ between seasons (*P*=0.562). A contrasting pattern of gametocyte carriage was observed. In the wet season, fewer infections harbored gametocytes (22.3% vs. 33.8%, *P*=0.009), but densities were 3-fold higher (*P*<0.001). Thus, in the wet season, among gametocyte positive individuals, higher proportion of all parasites were gametocytes, reflecting an increased investment in transmission.

## Introduction

Malaria control requires mapping potential silent gametocyte reservoirs in time and space. In many settings with pronounced seasonality in rainfall, *Anopheles* mosquitoes are few in the dry season as opposed to wet season where they are plentiful, resulting in transmission primarily occurring during and shortly after the wet season (Machani et al., 2020; Huestis & Lehmann, 2014; Jawara et al., 2008; Ouédraogo et al., 2008; Hamad et al., 2002). It is not known how far *Plasmodium falciparum* adapts its transmission potential to changes in vector abundance across seasons. Adaptions to increase transmission potential when chances for onward transmission are plenty could maximize the fitness of the parasite population. Understanding such adaptations are crucial to design transmission-reducing interventions.

Over the course of the red blood cell cycle, a small proportion of *P. falciparum* parasites develop into gametocytes, the sexual form of the parasite (Sinden, 1983). A mosquito blood meal needs to contain at least one female and one male gametocyte to be infective (Reece et al., 2008; Paul et al., 2000). The ingested gametocytes develop into oocysts and after approximately two weeks, into sporozoites that are transmitted to the next vertebrate host (Bruce et al., 1990). *P. falciparum* gametocytes exist in five morphologically distinguishable stages (Hawking et al., 1971). Early ring stage gametocytes circulate in peripheral blood (Farid et al., 2017) while late stages I-IV sequester for 7 to 12 days in inner organs including bone marrow and spleen until maturity (Farfour et al., 2012; Eichner et al., 2001; Paul et al., 2000). The mature stage V gametocytes re-enter the peripheral circulation where they require an additional 3 days to become fully infective (Lensen et al., 1999; Smalley & Sinden, 1977). Stage V gametocytes remain in the circulation for a mean period of 6.4 days to a maximum of 3 weeks (Eichner et al., 2001). Due to the sequestration of developing gametocytes, they are rarely detected in peripheral blood during the first two weeks following sporozoite inoculation.

A large proportion of all *P. falciparum* infections remain asymptomatic. Untreated infections can persist for several months (Rodriguez-Barraquer et al., 2018; Moormann et al., 2013; Nassir et al., 2005). During this time, parasite densities fluctuate and are often below the limit of detection by microscopy. Transmission stemming from asymptomatic infections is a key obstacle for malaria control and elimination. A previous study in western Kenya found asymptomatic individuals to be more infective than clinical cases (Gouagna et al., 2004). Even after antimalarial treatment, gametocytes may continue to circulate for up to 2-3 weeks (Bousema et al., 2010). Gametocyte densities are an important measure to predict the infectiousness of humans to mosquitoes (Gonçalves et al., 2017; Ouédraogo et al., 2016; Churcher et al., 2013), and thus useful for evaluating the effects of interventions that aim to reduce transmission (malERA, 2017).

Gametocyte density in the blood is governed by the conversion rate, i.e., the proportion of early ring stage parasites committed to sexual vs. asexual development. A higher proportion of parasites developing into gametocytes will increase transmission if vectors are present. On the other hand, the investment in gametocytes is lost if gametocytes are not taken up by mosquitoes. The factors affecting the conversion rate are not well understood. In laboratory culture and rodent malaria models, factors such as high parasite density (Mitri et al., 2009) and drug pressure (Buckling et al., 1999) have been found to impact gametocyte conversion. Few studies have measured the conversion rate directly in natural infections and observed pronounced variation (Usui et al., 2019; Poran et al., 2017; Smalley et al., 1981).

Areas of western Kenya experience perennial malaria transmission with peaks in vector density and transmission coinciding with seasonal rains in April-August and October-November (Machani et al., 2020; Desai et al., 2014). In regions with pronounced seasonality in vector abundance, parasites could increase their fitness by increasing their gametocyte conversion rate in the wet season. A small study involving 25 individuals in Sudan observed such a pattern (Gadalla et al., 2016). However, it remains unclear whether this is a general phenomenon, i.e., whether asymptomatic *P. falciparum* infections modulate the investment in gametocytes to coincide with the appearance of vectors at the start of transmission period.

To understand seasonal changes in gametocyte carriage, we compared *P. falciparum* gametocyte densities in asymptomatic individuals between the dry and wet seasons in a low-transmission setting (Homa Bay) and a moderate-transmission setting (Chulaimbo) in western Kenya. Blood stage parasites were diagnosed by *var*ATS qPCR, and mature female gametocytes were quantified using *pfs25* reverse transcriptase qPCR.

## Results

### Prevalence and density of *P. falciparum* infections

2859 samples with age distribution representative of the population were analyzed in this study. The demographic characteristics of the study participants are summarized in Table 1.

**Table 1.**
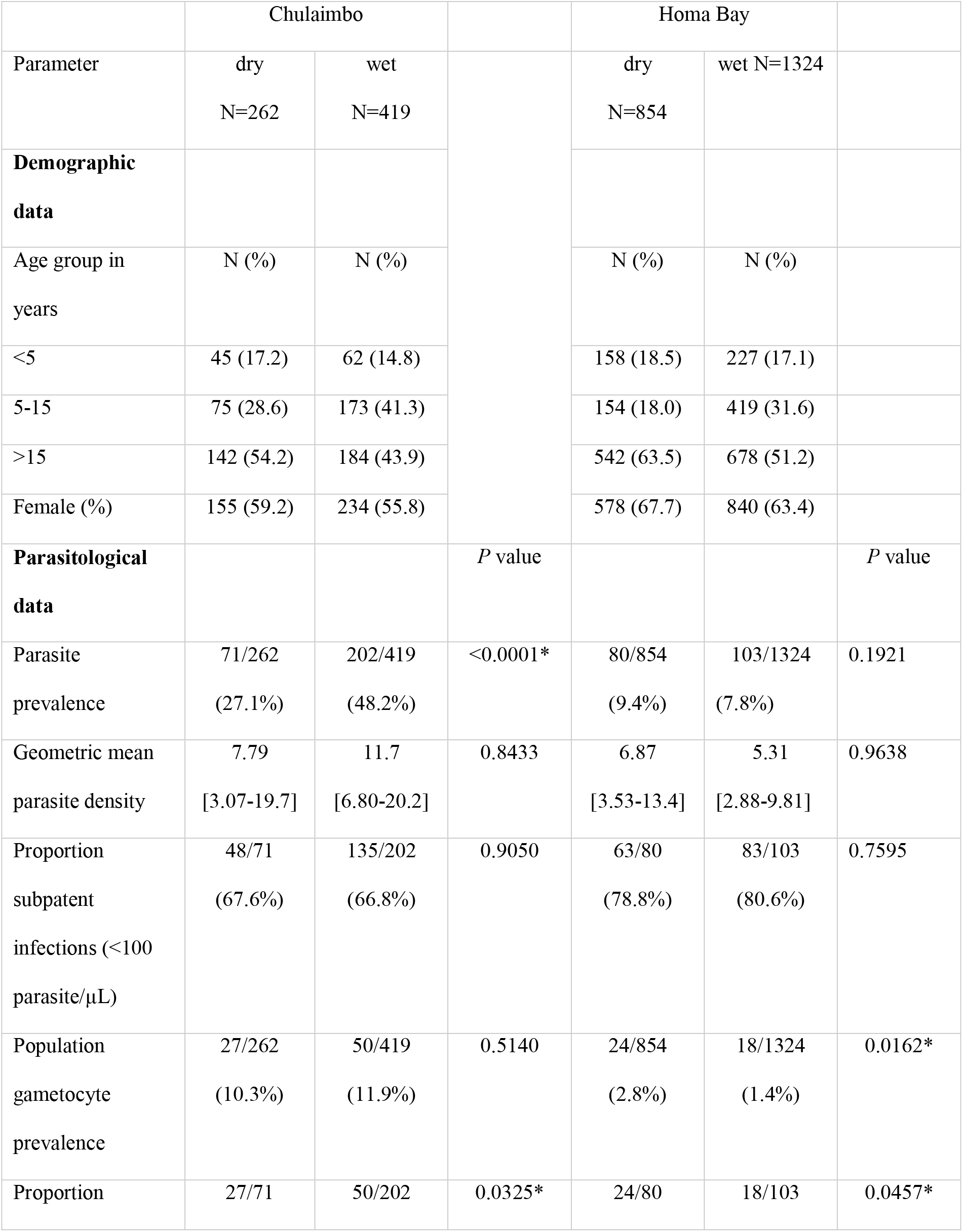
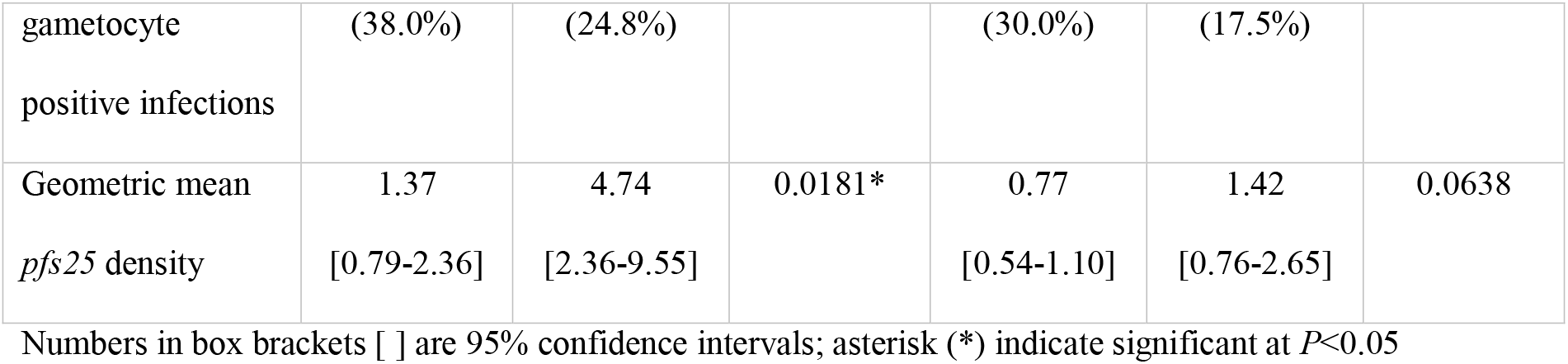
Characteristics of study participants

In both seasons, prevalence of *P. falciparum* infection was significantly higher in Chulaimbo than Homa Bay (wet: *P*<0.001, dry: *P*<0.001, Table 1). In Chulaimbo, the prevalence was significantly higher in the wet season (*P*<0.001, Table 1), but did not differ between seasons in Homa Bay (*P*=0.192, Table 1). Across all surveys, prevalence was higher in males than females (21.4% vs. 12.8%, *P*<0.001). School-age children (5-15 years) were at highest risk of infection (Fig 1).

**Figure 1.**
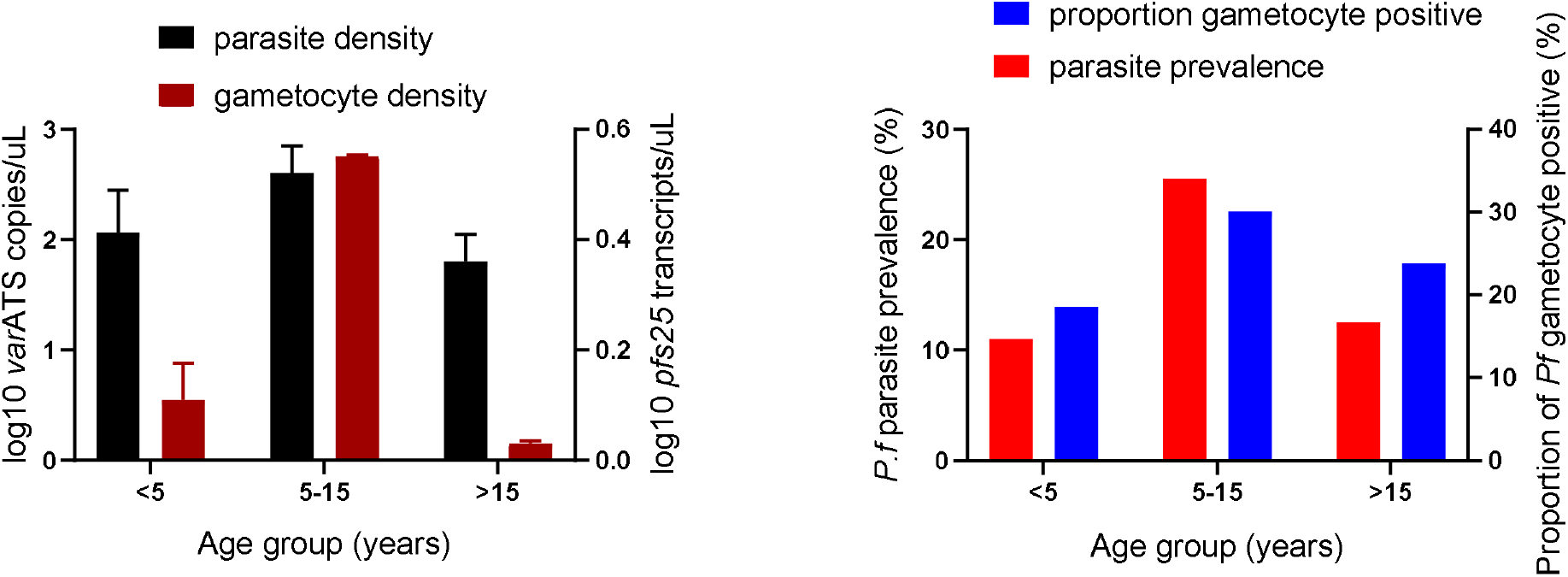
Age trends in *P. falciparum* parasite and gametocyte prevalence and density. Blood stage parasite density was measured by *var*ATS qPCR, and gametocyte density by *pfs25* mRNA reverse transcription qPCR. The proportion of gametocyte positive infections refers to proportion of all individuals with blood stage parasite who were positive for gametocytes.

Parasite densities by qPCR did not differ between seasons (Table 1). Across all surveys, parasite densities differed significantly between age groups (*P*<0.001, Fig 1). The densities peaked in children aged 5-15 years with a mean of 20.8 parasites/μL (95% confidence interval [CI95]: 12.6-34.5), and thus were 6-fold higher than in adults aged >15 years (3.3 parasites/μL, CI95: 2.1-5.1).

### Proportion of gametocyte positive infections and gametocyte density

Across all surveys, gametocytes were detected in 119/2859 (4.2%) individuals. The population gametocyte prevalence differed significantly between sites across seasons (Table 1, wet: *P*<0.001, dry: *P*<0.001), and ranged from 1.4% in Homa Bay in the wet season to 11.4% in Chulaimbo in the wet season (*P*<0.001, Table 1). The proportion of all individuals with blood stage parasites who were positive for gametocytes (the proportion of gametocyte positive infections) was significantly higher in the dry season (33.2%) than in wet season (22.3%, *P*=0.009, Table 1), but no difference was observed between sites (wet: *P*=0.149, dry: *P*=0.298).

The proportion of parasite and gametocyte carriers, and proportion of gametocyte positive infections was highest in school-age children aged 5-15 years across seasons and sites (Fig 1). *pfs25* transcripts/μL differed significantly between age groups (*P*=0.004, Fig 1). The transcript copies/μL peaked in children aged 5-15 years with a mean of 3.6 transcript/μL (CI95: 2.0-6.4), and thus were 3-fold higher than in adults aged >15 years (1.1 transcript/μL, CI95: 0.8-1.5). The correlation between *varATS* copy numbers and *pfs25* transcripts was moderate, but highly significant (R=0.36, *P*<0.001, Fig 2). Likewise, the probability to detect gametocytes was correlated with parasite density. Each 10-fold increase in genome copies resulted in 3.23-fold higher odds in carrying *pfs25* transcripts.

**Figure 2.**
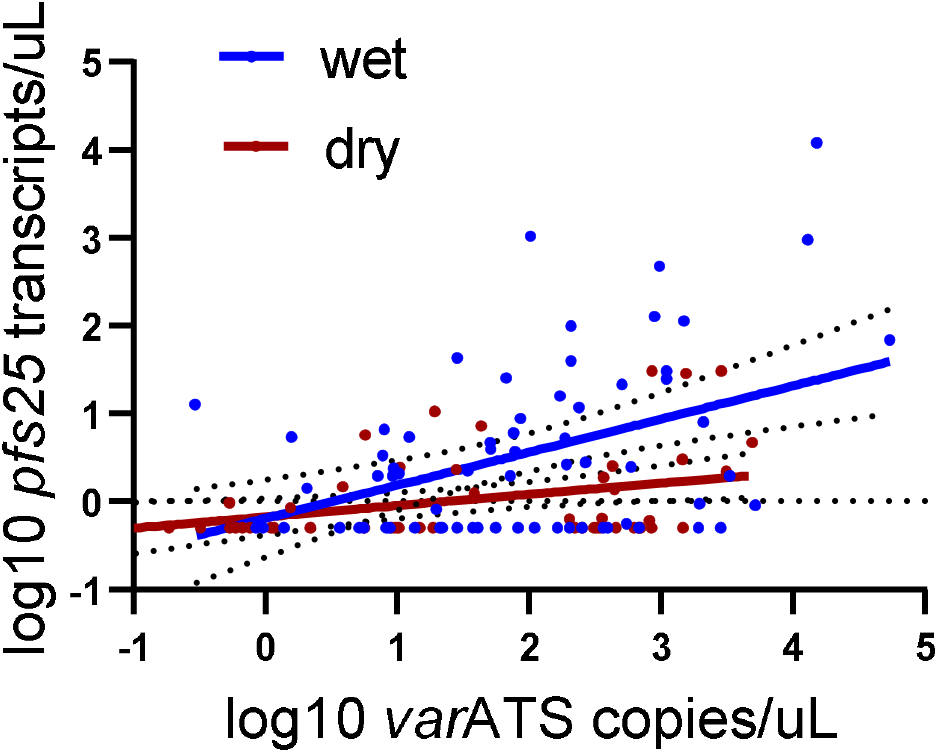
Correlation between *P. falciparum* asexual parasite and gametocyte densities across seasons. Dotted lines show 95% confidence intervals.

### Seasonal differences in gametocyte carriage

Seasonal patterns in gametocyte carriage were similar in Homa Bay and Chulaimbo. Thus, results are presented for both sites combined, with site-specific data in Table 1. The proportion of gametocyte positive infections was significantly higher in the dry season (33.8%, 51/151) compared to the wet season (22.3%, 68/305, *P*=0.009). In contrast, mean gametocyte densities were 3-fold higher in the wet season (wet: 3.46 *pfs25* transcripts/μL (CI95: 2.0-6.0), dry: 1.05 *pfs25* transcripts/μL (CI95: 0.8-1.5), *P*<0.001, even though parasite densities did not differ across seasons (wet: 8.98 *var*ATS copies/genome (CI95: 5.9-13.6), dry: 7.29 *var*ATS copies/genome (CI95: 4.2-12.7), *P*=0.562). The difference in *pfs25* transcript numbers between seasons remained highly significant when including log-transformed parasite densities as a predictor in multivariable analysis (Table 2). No interaction was observed between parasite density and the probability that an individual carried gametocytes, and season (*P*=0.739).

**Table 2.**
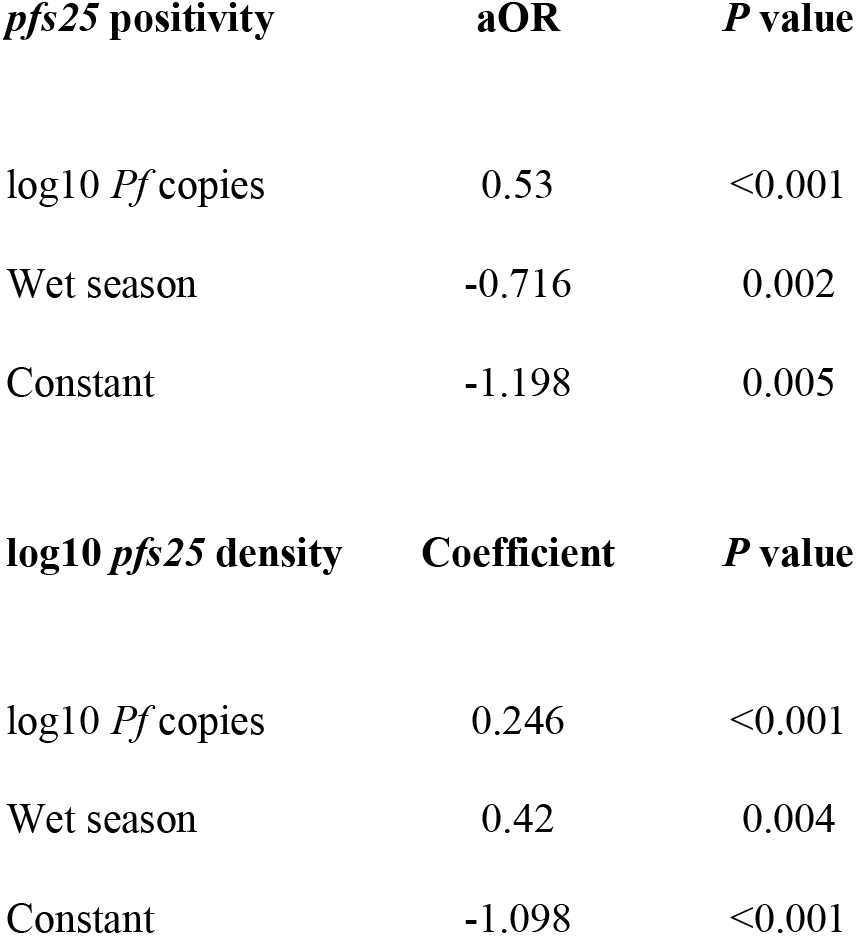
Multivariable predictors of gametocyte positivity and density

| *pfs25* positivity | aOR | *P* value |
| --- | --- | --- |
| log<sub>10</sub> *Pf* copies | 0.53 | <0.001 |
| Wet season | -0.716 | 0.002 |
| Constant | -1.198 | 0.005 |

| log<sub>10</sub> *pfs25* density | Coefficient | *P* value |
| --- | --- | --- |
| log<sub>10</sub> *Pf* copies | 0.246 | <0.001 |
| Wet season | 0.42 | 0.004 |
| Constant | -1.098 | <0.001 |

Pronounced variation in gametocyte carriage was observed among infections, with many medium- or high-density infections not carrying any detectable gametocytes. Among infections with a density of >2000 *var*ATS copies/μL (corresponding to >100 parasites/μL), in the wet season 60.0% (24/40) carried gametocytes versus 36.8% (32/87) in the dry season (*P*=0.014). Very few individuals carried gametocytes at high densities. For example, across both surveys, only 30 individuals carried *pfs25* transcripts at densities >5 transcripts/μL, 6 in the dry and 24 in the wet season. Among them, 4/6 and 19/24 were school-age children aged 5-15 years.

In multivariable analysis, only parasite density and season where found to be significantly associated with the probability that an individual was gametocyte positive (Table 2). Age group (*P*=0.195), sex (*P*=0.214), and site (*P*=0.364) were not associated. Likewise, gametocyte density was only significantly associated with parasite density and season, but not site (*P*=0.063), age group (*P*=0.733), or sex (*P*=0.611) (Table 2).

### Gametocyte carriage among patent and subpatent infections

A sensitivity of 100 blood stage parasites/μL (i.e. asexual parasites and gametocytes) was assumed to determine the proportion of infections that could be detected by Rapid Diagnostic Test (RDT) or light microscopy. Given this threshold, 72.1% (329/456) of all infections were subpatent across all surveys. No difference in the proportion of subpatent infections was observed between seasons (dry: 73.5% (111/151), wet: 71.5% (218/305), *P*=0.648). 52.9% (63/119) of all infections with gametocytes detected by RT-qPCR were subpatent across all surveys with equal proportions in the dry (52.9% (27/51)) and wet season (52.9% (36/68)). Mean *pfs25* densities were 3-fold lower in subpatent infections compared to patent infections (1.26 vs. 3.64 transcripts/μL, *P*=0.003).

## Discussion

We observed a contrasting pattern of gametocyte carriage between the dry and the wet season in blood samples collected from 2859 healthy afebrile individuals residing in a malaria endemic area of western Kenya. In the wet season, when most transmission is expected to occur, fewer infections harbored gametocytes, but gametocyte densities were higher. The higher gametocyte densities in the wet season are particularly noteworthy as parasite densities did not differ between seasons. Thus, the proportion of gametocytes among total blood stage parasites was higher in the wet season compared to the dry season. Our results imply that parasites increase investment in gametocytes in the high transmission period to be synchronized with increased vector abundance in the rainy season. However, the adjustment was not uniform across all infections. Less than a quarter of infections carried detectable gametocytes in the wet season. Among low-density asymptomatic infections gametocytes might be below the limit of detection even by RT-qPCR (Koepfli & Yan, 2018). Yet, in the current study, even among medium-to-high density infections (above 100 parasites/μL), more than half did not carry gametocytes. Given the high sensitivity of our RT-qPCR, limited detectability cannot cause this result.

While our quantification of *pfs25* transcripts is a good marker of infectivity at time of sample collection (Bradley et al., 2018; Gonçalves et al., 2017; Churcher et al., 2013), it is only an indirect measure of commitment to transmission. Asexual parasite densities are expected to peak early in the infection, when mature gametocytes are not yet circulating. Likely, some of the high-density infections were recently acquired and carried sequestered gametocytes that appeared in the blood a few days after sample collection. Among infections with above average proportions of gametocytes, asexual densities might have been higher two weeks prior when gametocyte development was initiated. Alternatively, the pattern might reflect true differences in gametocyte conversion. Few studies have measured the conversion rate directly on field isolates, and found pronounced variation among *P. falciparum* strains (Usui et al., 2019; Poran et al., 2017; Smalley et al., 1981). The factors underlying these differences remain poorly understood.

Our findings of higher gametocyte densities in the wet season are in line with xenodiagnostic surveys conducted from asymptomatic residents of Burkina Faso and Kilifi, Kenya. Gametocyte densities determined by molecular assays targeting *pfs25* transcripts and infectivity were substantially higher in the wet compared to the dry season (Gonçalves et al., 2017; Ouédraogo et al., 2016). Similarly, the present study corroborates previous work on asymptomatic individuals in eastern Sudan, in which gametocyte densities significantly increased during the period of expected mosquitoes appearance relative to the transmission-free season with no corresponding substantial increase in parasite densities (Gadalla et al., 2016). Increasing investment in gametocytes is beneficial to the parasite in maximizing onward transmission when mosquitoes are plentiful.

As opposed to Chulaimbo where parasite prevalence doubled in the wet season, in Homa Bay the prevalence did not change. The variations in seasonal parasite prevalence pattern between Chulaimbo and Homa Bay may be due to differences in species composition of local vector populations (Ayanful-Torgby et al., 2018). In Chulaimbo, *An. Arabiensis* forms the predominant mosquito vector species followed by *Anopheles gambiae s.s.* (Machani et al., 2020), whereas in Homa Bay *An. funestus* is the predominant mosquito vector species (McCann et al., 2014). *An. funestus* prefers permanent bodies of water like irrigated rice fields that last beyond the wet seasons, while *An. arabiensis* prefers temporary holes and pools that dry out once the rainy season ends (Kweka et al., 2012; Mala & Irungu, 2011; Ndenga et al., 2011; Fillinger et al., 2004).

In all surveys, 67-80% of infections were subpatent. In both sites and seasons, approximately half of all individuals that had gametocyte detected by RT-qPCR carried infections at densities below the limit of detection of microscopy or rapid diagnostic test. They thus would escape screening of asymptomatic individuals using field-deployable diagnostics. Gametocyte densities were 3-fold lower in subpatent individuals, thus they would likely infect fewer mosquitoes than patent individuals. Subpatent *P. falciparum* gametocyte carriers in natural infections have the potential to infect mosquitoes (Gonçalves et al., 2017; Ouédraogo et al., 2016; Churcher et al., 2013). However, the contribution of these infections to transmission in different settings is not known (malERA, 2017).

*P. falciparum* parasite prevalence and density, and gametocyte prevalence and the proportion of gametocyte-positive infections were highest in school-age children. The higher mean gametocyte density in school-age children mirrored asexual parasite densities and no impact of host age on gametocyte carriage independently of parasite density was apparent. Yet, among the small number of individuals carrying gametocytes at moderate-to-high densities (>5 transcripts/uL), three quarters were in this age group. Our findings are in line with several studies that had identified this age group as an important source of ongoing malaria transmission (Coalson et al., 2018; Gonçalves et al., 2017; Coalson et al., 2016; Ouédraogo et al., 2016; Churcher et al., 2013).

## Conclusions

We have observed changes in the investment in transmission across seasons in a large survey of asymptomatic *P. falciparum* infections. Future research is needed to investigate how parasites sense changes in seasonality and to understand the factors underlying the increase in gametocyte density in the wet season. Our findings confirm that seasonality is an important aspect to consider when designing control measures targeted at asymptomatic carriers. The frequent carriage of gametocytes in the dry season implies that these infections constitute an important reservoir that initiate transmission in the wet season. A small number of individuals, mostly school children, carried very high gametocyte densities and likely contributed disproportionally to transmission. Targeted treatment of school children at the beginning of the wet season might thus reduce transmission substantially.

## Materials and Methods

### Study sites and participants

2859 asymptomatic individuals were sampled in cross-sectional surveys in the dry season (n=1116) between January and March 2019, and the wet season (n=1743) between June and August 2019 in Western Kenya, in Homa Bay (low transmission) and Chulaimbo (moderate transmission) (Table 1, Figure 3). In these areas, *P. falciparum* is the primary malaria parasite species (Idris et al., 2016). The study population included asymptomatic individuals aged 2 months to 99 years with no clinical symptoms. None of the study participants had been treated with antimalarial drugs within the three days prior blood sampling.

**Figure 3.**
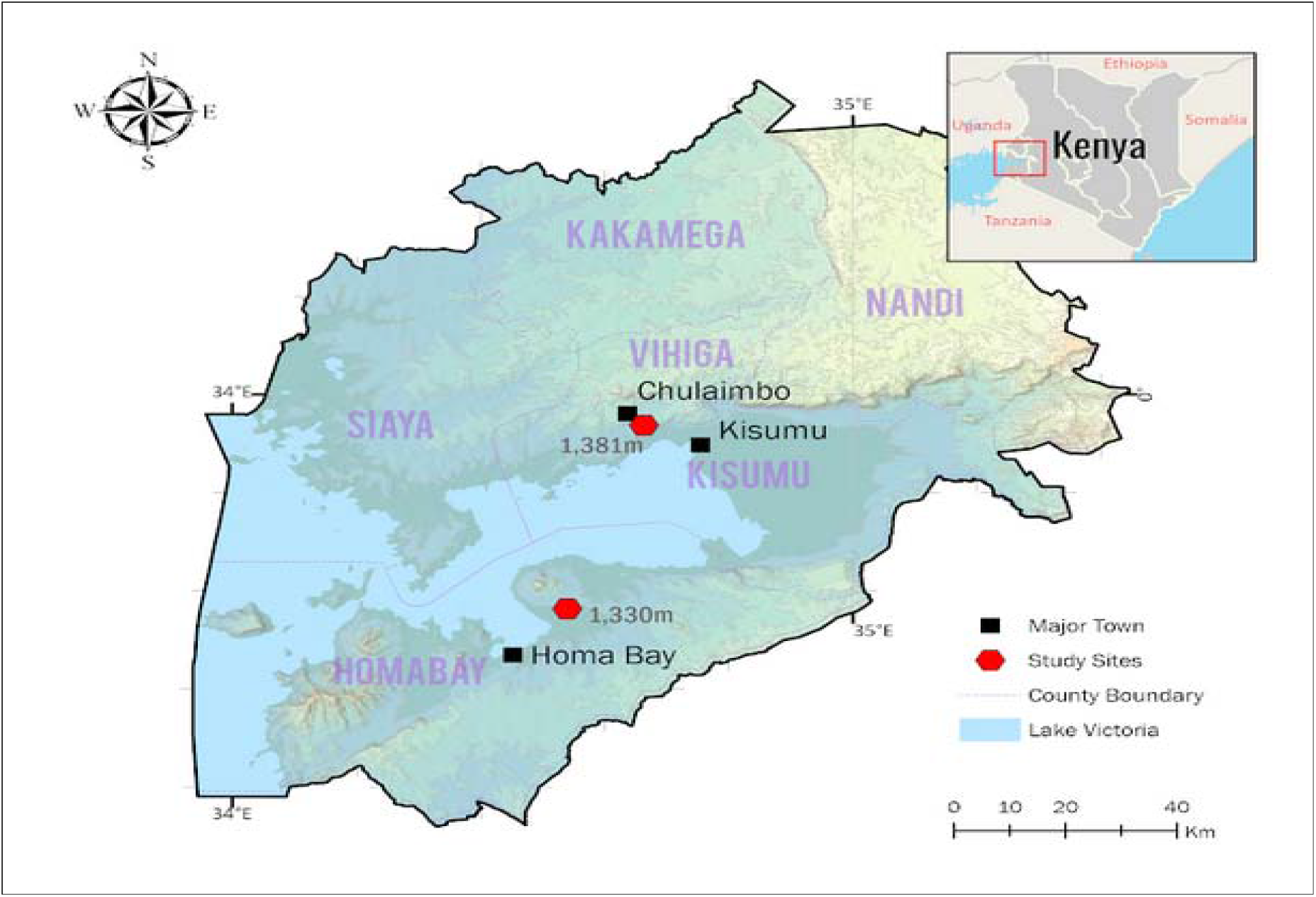
Map of study sites.

In Chulaimbo, *Anopheles arabiensis* is the primary vector. It is abundant in the wet season. *An. gambiae s.s* is the second predominant mosquito vector (Machani et al., 2020). In Homa Bay, *An. funestus* has re-emerged as the predominant species following development of pyrethroid resistance (McCann et al., 2014). According to the Kenya “End of Spray” Report (2018), indoor residual spraying (IRS) in Homa Bay has resulted in a reduction in malaria vector densities and sporozoite rates compared with Chulaimbo, where IRS has not been implemented.

### Ethical consideration

Ethical approval to conduct the study was obtained from Maseno University Ethics Review Committee (MUERC protocol number 00456), the University of California, Irvine Institutional Review Board (HS# 2017-3512), and the University of Notre Dame (#20076141). All study participants and guardians of minors gave informed consent prior to obtaining clinical and demographic information and drawing a blood sample.

### Sample collection and processing

350-400 μL of capillary blood was collected into EDTA microtainer tubes (Becton Dickinson, New Jersey, United States) by finger prick. For RNA preservation, 100 μL of whole blood was transferred to a tube containing 500 μL of RNAlater (Sigma-Aldrich, Missouri, United States) within 2 hours of collection and stored at −80°C until RNA extraction (Koepfli et al., 2015; Wampfler et al., 2013). The remaining blood was centrifuged, plasma removed and stored at −20°C. The red cell pellet was stored at −20°C until DNA extraction.

### Molecular parasite screening and quantification

DNA was extracted from 100 μL blood using the Genomic DNA Extraction kit (Macherey-Nagel, Düren, Germany) and eluted in an equivalent volume of elution buffer. DNA was screened for *P. falciparum* using ultrasensitive qPCR that amplifies a conserved region of the *var* gene acidic terminal sequence (*var*ATS) according to a previously published protocol (Hofmann et al., 2015). The *var*ATS gene assay amplifies ~20 copies/genome (Hofmann et al., 2015). The qPCR results were converted to *var*ATS copies/μL using external standard curve of ten-fold serial dilutions (5-steps) of 3D7 *P. falciparum* parasites quantified by droplet digital PCR (ddPCR) (Koepfli et al., 2016). The ddPCR thermocycling conditions, sequences and concentration of primers and probe are given in supplementary materials. Asexual parasite densities were calculated by dividing *var*ATS copy numbers by 20, reflecting the approximate number of copies per genome.

For all the gametocytes assays, RNA was extracted using the pathogen Nucleic Acid Extraction kit (Macherey-Nagel, Düren, Germany) and eluted in 50 μL elution buffer, i.e., RNA was concentrated two-fold during extraction. RNA samples were DNase treated (Macherey-Nagel, Düren, Germany) to remove genomic DNA that could result in a false positive *pfs25* signal (Meerstein-Kessel *et al*., 2018). A subset of RNA samples was tested by varATS qPCR, and all tested negative.

### Molecular gametocyte screening and quantification

For gametocyte detection by reverse-transcription quantitative PCR (RT-qPCR), RNA was extracted from all *P. falciparum* qPCR-positive samples. Gametocytes were quantified by the female *pfs25* mRNA transcripts using one-step RT-qPCR assays (Alkali Scientific, Florida, United States). All qPCR conditions, sequences and concentration of primers and probes are given in supplementary materials. The *pfs25* RT-qPCR results were converted to *pfs25* transcript copies/μL using external standard curve of ten-fold serial dilutions (5-steps) of 3D7 culture parasites quantified by ddPCR (supplementary materials).

### Statistical analysis

Parasite and gametocyte densities were log_10_ transformed and geometric means per μL blood calculated whenever densities were reported. The Shapiro-Wilk test and graphical normality was employed to determine normal distribution of data following log transformation. Differences in prevalence between seasons and sites were determined using the χ^2^ test. Differences in densities between seasons and sites were determined using T-test. Differences in densities between age-groups were determined by ANOVA’s Tukey’s multiple comparisons test. Multivariable analysis was employed to determine association of age, site and season with asexual parasite and gametocyte positivity and density. The associations were investigated by regression analysis. Pearson’s correlation test was conducted to establish the relationship between asexual parasite and gametocyte densities. Data analysis was done in GraphPad Prism version 8 and STATA version 14.

## Abbreviations

IRS: indoor residual spraying
RT-qPCR: reverse transcriptase – quantitative polymerase chain reaction
ddPCR: droplet digital PCR
*var*ATS: *var* gene acidic terminal sequence
RNA: ribonucleic acid
DNA: deoxyribonucleic acid
EDTA: ethylenediaminetetraacetic acid

## Author contributions

COO led sample collection, conducted lab work and primary data analysis, and wrote and revised the manuscript. SO, BNO, HA, and AKG helped with conceptualization, project administration, and supervised field work, M-CL supported project administration and data analysis, AED helped with conceptualization and project administration, JWK and GY conceived the study, acquired the funding, and supported project administration, CK supported conceptualization, and supervised lab work, data analysis, and manuscript writing and revision. All authors have read and agreed with the final version of the manuscript.

## Acknowledgements

We are grateful to all study participants. We thank the field teams of the sub-Saharan Africa ICEMR for support with sample collection. This work is published with the permission of the Director of KEMRI. This work was supported by grants from the National Institutes of Health (U19 AI129326, D43 TW001505, R01 AI050243, R01 AI130131). The funders had no role in study design, data collection and analysis, decision to publish or preparation of the manuscript.

## Supplementary materials

### Droplet digital PCR *var* ATS protocol

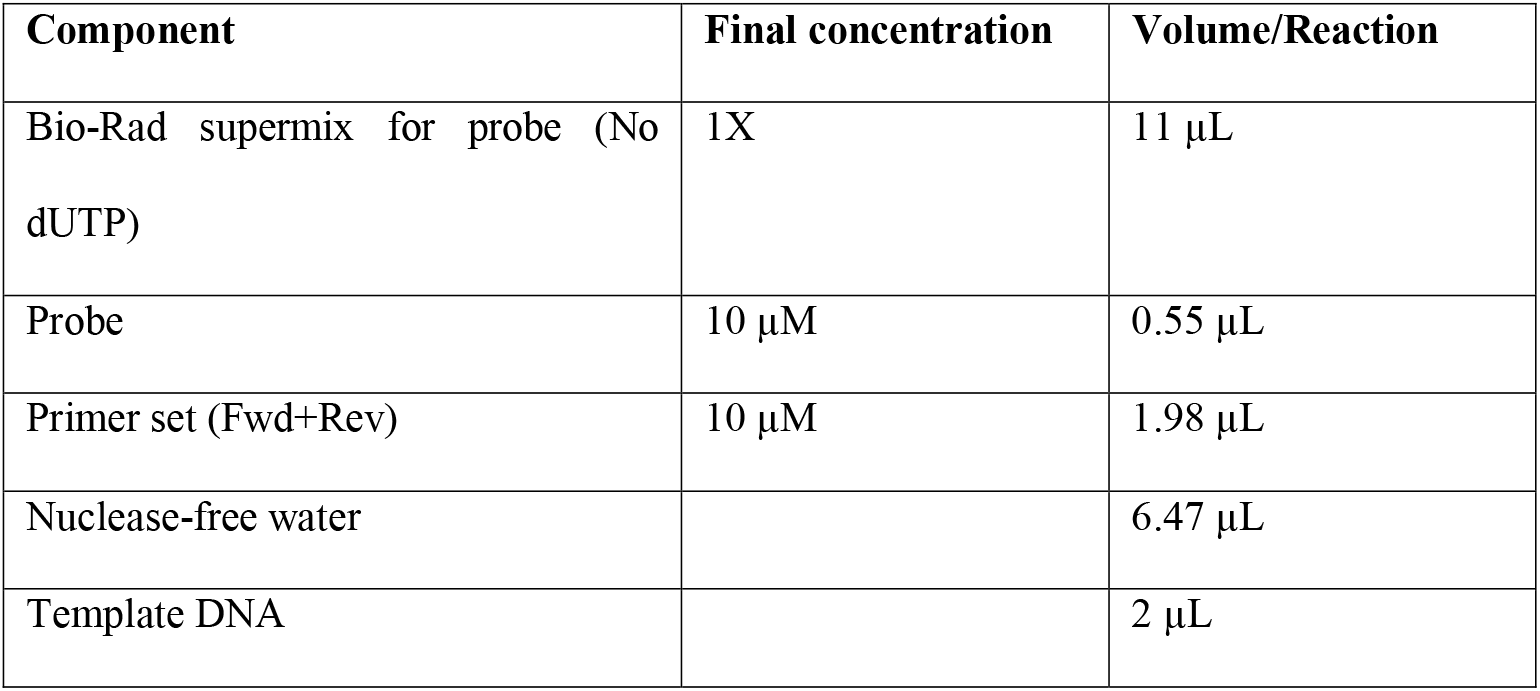

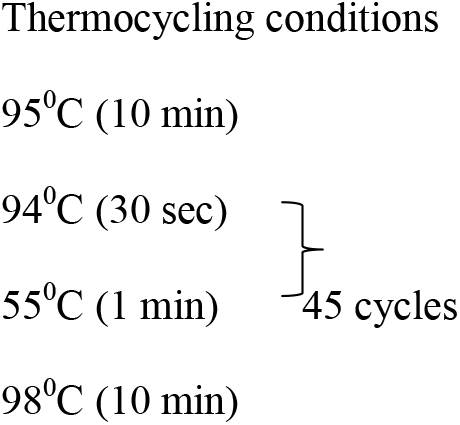

Primers and probe

*var*ATS fwd CCCATACACAACCAAYTGGA
*var*ATS rev TTCGCACATATCTCTATGTCTATCT
*var*ATS probe 6-FAM-TRTTCCATAAATGGT-NFQ-MGB

### *pfs25* RT-qPCR gametocyte screening

Master mix (12 μL)

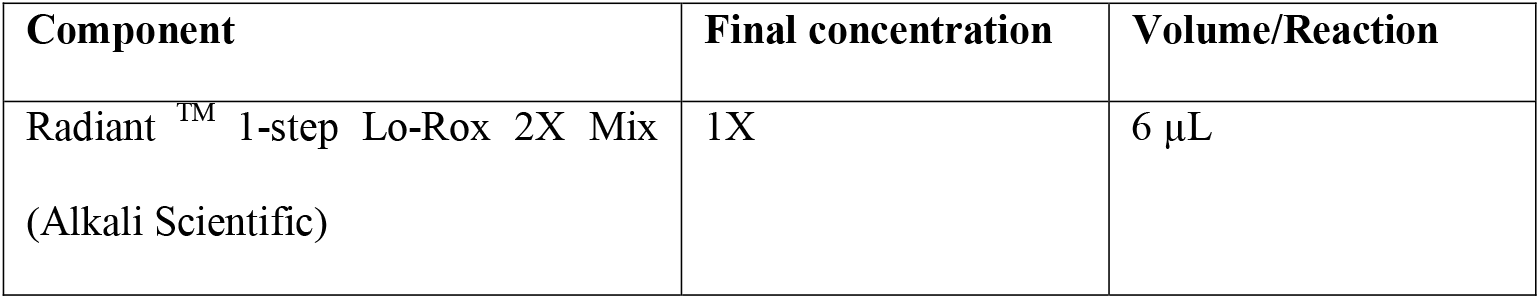
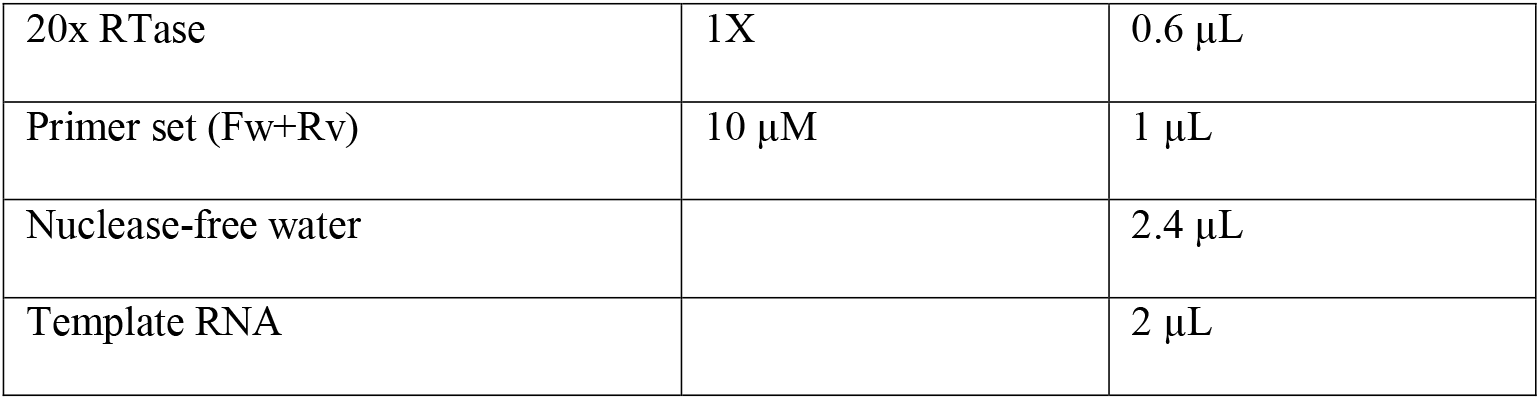

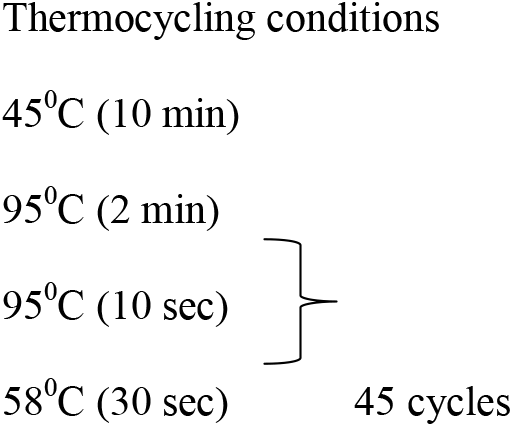

Primers

*pfs25* fwd CGT TTC ATA CGC TTG TAA ATG
*pfs25*_rev TTA ACA GGA TTG CTT GTA TCT AA

