## Supplementary Files for "*Plasmodium falciparum* increases its investment in gametocytes in the wet season in asymptomatic individuals"

**Supplementary materials**

**Droplet digital PCR *var*ATS protocol**

| **Component** | **Final concentration** | **Volume/Reaction** |
| --- | --- | --- |
| Bio-Rad supermix for probe (No dUTP) | 1X | 11 µL |
| Probe | 10 µM | 0.55 µL |
| Primer set (Fwd+Rev) | 10 µM | 1.98 µL |
| Nuclease-free water |  | 6.47 µL |
| Template DNA |  | 2 µL |

Thermocycling conditions

95^0^C (10 min)

94^0^C (30 sec)

55^0^C (1 min)	45 cycles

98^0^C (10 min)

Primers and probe

*var*ATS fwd	CCCATACACAACCAAYTGGA

*var*ATS rev	TTCGCACATATCTCTATGTCTATCT

*var*ATS probe	6-FAM-TRTTCCATAAATGGT-NFQ-MGB

***pfs25* RT-qPCR gametocyte screening**

Master mix (12 µL)

| **Component** | **Final concentration** | **Volume/Reaction** |
| --- | --- | --- |
| Radiant ^TM^ 1-step Lo-Rox 2X Mix (Alkali Scientific) | 1X | 6 µL |
| 20x RTase | 1X | 0.6 µL |
| Primer set (Fw+Rv) | 10 µM | 1 µL |
| Nuclease-free water |  | 2.4 µL |
| Template RNA |  | 2 µL |

Thermocycling conditions

45^0^C (10 min)

95^0^C (2 min)

95^0^C (10 sec)

58^0^C (30 sec)	45 cycles

Primers

*pfs25* fwd	CGT TTC ATA CGC TTG TAA ATG

*pfs25*_rev	TTA ACA GGA TTG CTT GTA TCT AA
